# The worldwide allometric relationship in anatomical structures for plant roots

**DOI:** 10.1101/2022.11.29.518307

**Authors:** Yue Zhang, Jing-Jing Cao, Qing-Pei Yang, Ming-Zuo Wu, Yong Zhao, De-Liang Kong

## Abstract

The anatomical structures, i.e., the cortex and stele, are fundamental for the absorptive function of plant roots. Unraveling how the allometric structures are assembled in absorptive roots is essential for our understanding the plant ecology, physiology and responses to global environmental changes. In this review study, we first compile a globally largest dataset on key root structural traits, i.e., root diameter, cortex thickness and stele radius across 512 species. Using this largest dataset, we confirm an allometric relationship of absorptive root structures in a previous study using a much smaller species pool, i.e., the cortex thickness increased much faster than the stele radius with increasing root diameter. The allometric relationship is further validated within and across different plant growth forms (woody, grass, and liana species), mycorrhiza types (arbuscular mycorrhiza, ectomycorrhiza, and orchid mycorrhizas), phylogenetic gradients (from ferns to Orchidaceae of primitive angiosperms), and environmental change scenarios (e.g., the elevation of atmospheric CO_2_ concentration and nitrogen fertilization), supporting the universal allometric relationship in plant roots. We then summarized recent proceedings as well as possible issues on mechanisms underlying the root allometric relationship. The ecological and evolutionary implications for this allometric relationship in roots are also discussed. Finally, we propose several directions that should be stressed in future studies regarding the allometric relationship in plant roots.

## 1. Introduction

Plant roots play a crucial role in plant growth, vegetation dynamics, ecosystem functioning like productivity formation, nutrient cycle and their responses to environmental changes (Carmona et al., 2021; Laughlin et al., 2021; Chandregowda et al., 2022; Encinas-Valero et al., 2022; Hong et al., 2022). Compared to studies on plant above-ground organs such as leaves and stems that have undergone enormous proceedings (Wright et al., 2004; Diaz et al., 2016; Joswig et al., 2022), our understanding of plant roots remains in its infancy. The core function of plant roots is to absorb soil water and nutrients, which is undertaken by a few terminal root branch orders, i.e., absorptive roots, mainly bearing primary root tissues (Guo et al., 2008b). Generally, the absorptive function is depicted by a range of traits in root morphology, physiology, anatomy, chemistry, mechanics and microbial symbiosis (McCormack et al., 2017; Wambsganss et al., 2021; Wen et al., 2022; Yan et al., 2022). Among these root traits, root diameter seems like the most important one given that it is closely associated with a suite of other root traits as well as mycorrhizal fungi apart from its well-known feature of easily measured and great inter-specific variation (Eissenstat et al., 2015; Li et al., 2018; Bergmann et al., 2020; Wen et al., 2022). Furthermore, root diameter is the phylogenetically most conservative root trait, suggesting that the great inter-specific variation could largely be an evolutionarily imprint from the geological environmental change such as atmospheric CO_2_ decline since the Cretaceous (Comas et al., 2012; Chen et al., 2013; Pineiro et al., 2020; Lugli et al., 2021).

The absorption function of plant roots is essentially determined by root anatomical structures. Generally, absorptive roots are composed of two cylindrical components, i.e., cortex and stele. The cortex directly takes part in the absorption of water and nutrients and indirectly acquire these resources by associations with mycorrhizal fungi (Brundrett, 2002; Ma et al., 2018; Rich et al., 2021). The stele is responsible for transporting water and nutrients upward to stems and leaves, and the stele supply energy demanding of the roots with leaf photosynthate. Theoretically, the change in root diameter is mainly derived from the size variations of the cortex and stele, while how the cortex and stele are coordinated with the shifts of root diameter has not been uncovered until 2014 when two research groups independently found an allometric relationship between the cortex and stele (Gu et al., 2014; Kong et al., 2014), i.e., the cortex thickness increased linearly and much faster than the stele radius with increasing root diameter. Later, this allometric relationship was reported in many studies, and was synthesized by Kong et al. (2019) who reported a global existence of the allometric relationship in root cortex and stele across 204 plant species.

Uncovering of the allometry relationship paves a new way for our understanding of the form-function relationship in roots and plant evolution and adaptation to environmental changes (McCormack et al., 2017; Kong et al., 2019; Bergmann et al., 2020; Zhou et al., 2022). Since the global recognition of the allometric relationship between root cortex and stele in 2019, mounting studies have focused on root anatomical traits for one hand, and for the other hand we also note that some early studies on Orchidaceae and vine root anatomy were neglected in Kong et al. (2019). Further, despite the worldwide mycorrhizal associations in terrestrial plants, we know little about whether and how the allometric relationship in absorptive roots varies among mycorrhizal types with contrasting mycorrhizal structures and functioning (e.g., arbuscular mycorrhiza (AM) *vs*. ectomycorrhiza (EM)) (Brundrett, 2002; Martin et al., 2017) Additionally, no studies to date have explored how the environmental changes affect the above allometric relationship in roots. Therefore, it is necessary to further test the generality of the root allometric relationship using a much larger species pool than that in Kong et al. (2019).

To fulfill this purpose, in this review paper, we made a thorough screening of the data on the cortex thickness and the stele radius of absorptive roots in Web of Science, Google Scholar, FRED 3.0 and CNKI (China’s national knowledge infrastructure) using keywords included “cortex”, “stele”, “anatomic structure”, “allometric relationship”, “root diameter”. Our searching yielded 3,676,679 papers and reports. We then refined these results according to additional criteria: (1) the study must be an empirical rather than a review or perspective; (2) the data on root diameter and stele radius were accessible. We also included some unpublished data (supplementary dada 2) on root anatomical traits which were measured at the same sites and following the same procedures as our previous study (Kong et al., 2014). Finally, our dataset included 32 empirical studies (supplementary data 1) with a total of 698 observations of 512 species at 41 sites (Fig. 1). Specifically, the dataset included 271 woody species and 241 non-woody species (78 grass, 92 herb, 37 fern and 28 Orchidaceae species). In addition, 13 liana species were included in the dataset. For the same species appearing in different studies, we used the average value of the root traits across species as the trait value of this species. For some studies with only the data of the stele radius and root diameter, we use the difference between the root diameter and stele radius (equal to the thickness of tissues outside the stele including the epidermis, exodermis and cortex, i.e., tToS, as a proxy approximate to the cortex thickness (Kong et al., 2019). For the studies with root trait data displayed in figures or only photos on root anatomical structures presented, we digitalized the root trait data using the software “SigmaScan Pro software (V5.0, SPSS Inc., Chicago, USA)” and the software “IMAGE J (NIH Image, Bethesda, MD, USA)”, respectively.

**Fig. 1.**
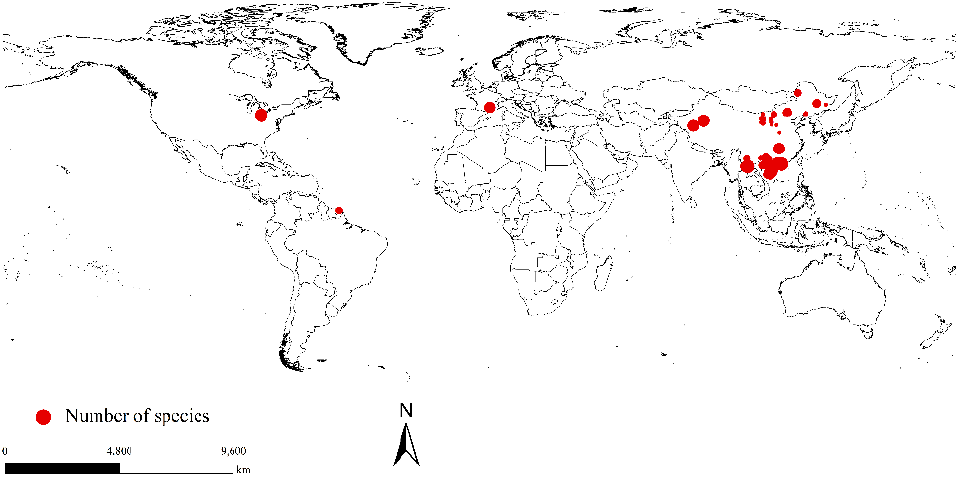
A global map of 32 studies reporting data of the cortex and the stele in absorptive roots. Each study is represented by a red circle, and the size of the circle is proportional to the species number included in the studies.

Overall, in this review we aim to: (1) test the generality of the allometric relationship between root cortex and stele within and across different plant growth forms, mycorrhizal types and environmental treatments; (2) summarize mechanisms and implications for such allometric relationship; (3) propose important directions for future studies regarding the root allometric relationship.

## 2. Generality of the allometric relationship in absorptive roots

The allometric relationship between the cortex and the stele still held across the 698 observations and 512 species of root anatomical structures (Table 1; Fig. 2a, 2b). For the 26 studies with root anatomical structures examined in at least three species, the allometric relationship in absorptive roots existed in most of the studies, while only four studies seemed exceptional (Table 2; Fig. 3; Supplementary data 1).

**Table 1.**
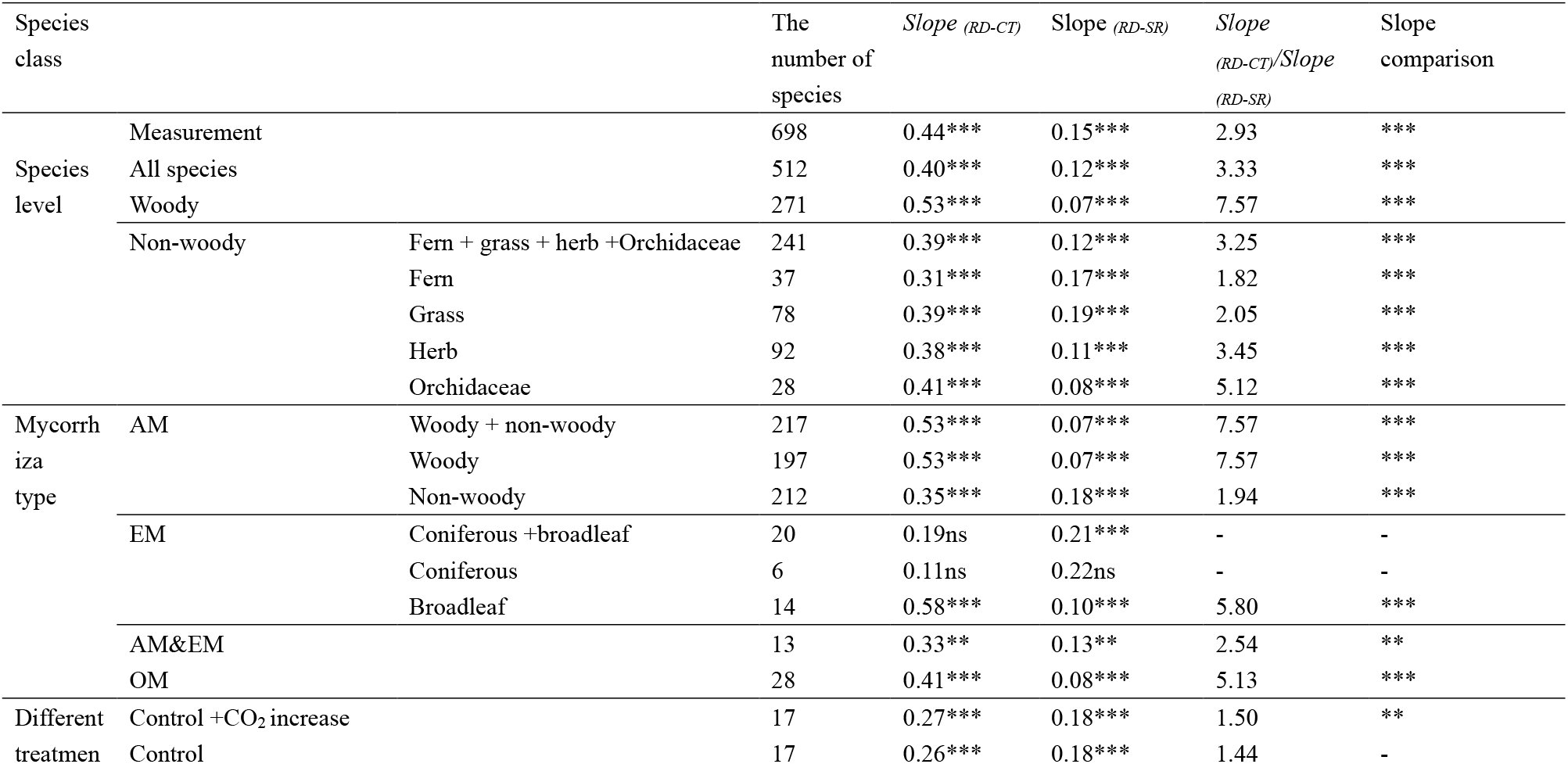
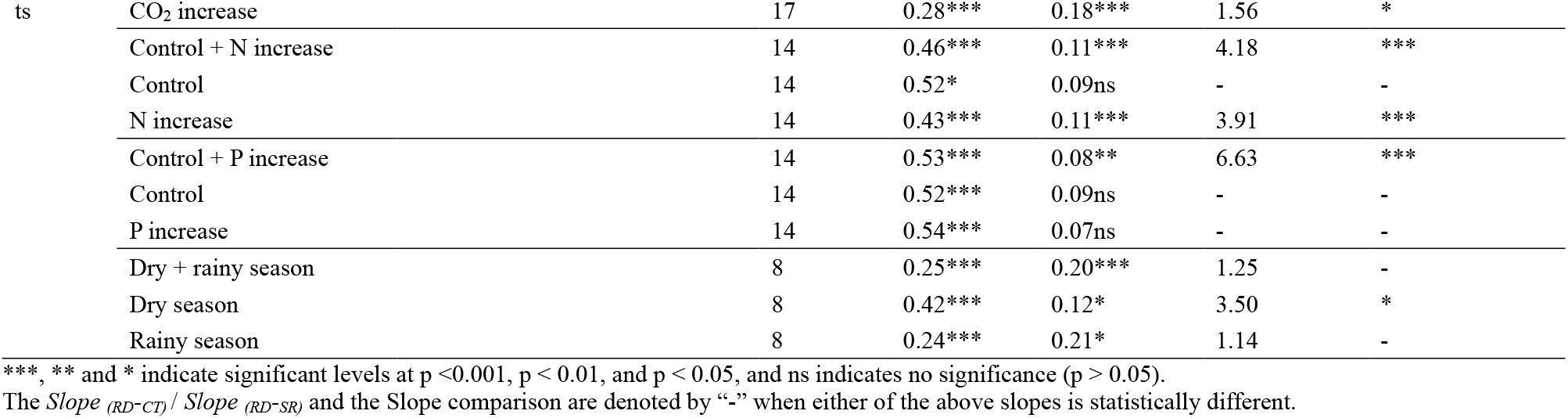
The relationships between root dimeter (RD) and the cortex thickness (CT) and the stele radius (SR) for different plant forms and different environmental treatments. *Slope _(RD-CT)_*: the slope for the regression of the cortex thickness with root diameter; *Slope _(RD-SR)_* the slope for the regression of the stele radius with root diameter; *Slope _(RD-CT)_* / *Slope _(RD–SR):_* ratio of the *Slope _(RD-CT)_* to the *Slope _(RD-SR)_;* Slope comparison: the statistical test of the difference between the above two slopes using standardized major axis method.

**Fig. 2.**
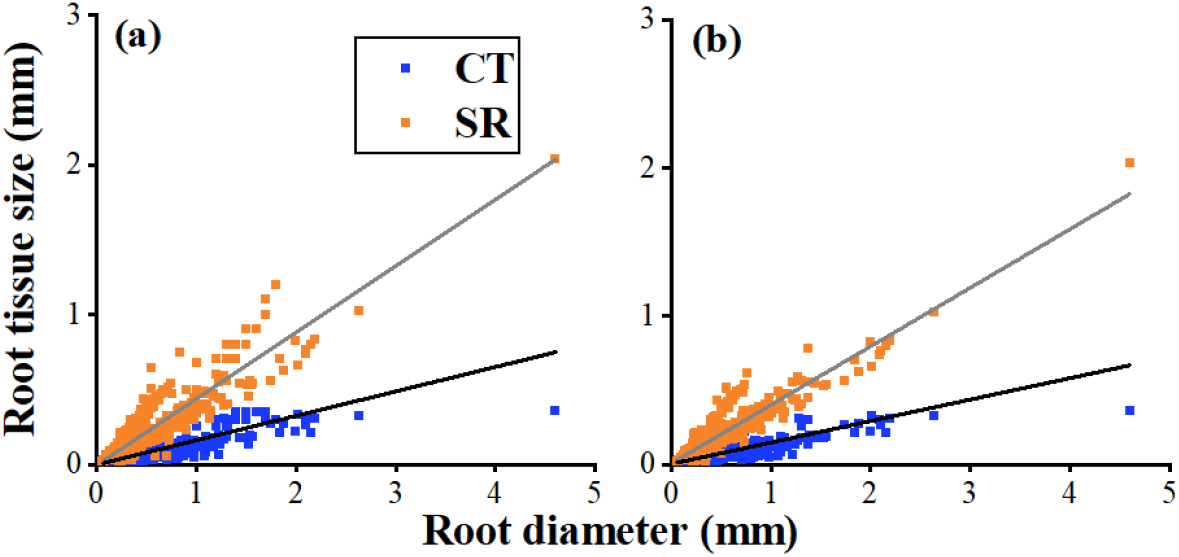
The Allometric relationship between the root cortex and the stele across 698 observation (a) and 512 species (b), CT: cortex thickness; SR: stele radius.

**Table 2.** The allometric relationships between root dimeter (RD) and the cortex thickness (CT) and the stele radius (SR) in each of the 26 studies with more than 3 plant species examined. See supplementary data 1 for the details of these studies. *Slope _(RD-CT)_*: the slope for the regression of the cortex thickness with root diameter; *Slope _(RD-SR)_*: the slope for the regression of the stele radius with root diameter; *Slope _(RD-CT)_/ Slope _(RD-SR):_* ratio of the *Slope _(RD-CT)_* to the *Slope _(RD-SR)_;* Slope comparison: the statistical test of the difference between the above two slopes using standardized major axis method.

| Study | The number of species | *Slope* <sub>(RD-CT)</sub> | *Slope* <sub>(RD-SR)</sub> | *Slope* <sub>(RD-CT)</sub>/*Slope* <sub>(RD-SR)</sub> | Slope comparison |
| --- | --- | --- | --- | --- | --- |
| 1 | 19 | 0.27*** | 0.23*** | 1.17 | - |
| 2 | 14 | 0.37*** | 0.13* | 2.85 | * |
| 3 | 20 | 0.39*** | 0.12*** | 3.25 | *** |
| 4 | 23 | 0.37*** | 0.14*** | 2.64 | ** |
| 5 | 27 | 0.40*** | 0.10*** | 4.00 | *** |
| 6 | 5 | 0.39** | 0.07* | 5.57 | ** |
| 7 | 96 | 0.75*** | 0.10*** | 7.50 | *** |
| 8 | 9 | 0.47*** | 0.03ns | - | - |
| 9 | 12 | 0.39*** | 0.11*** | 3.55 | *** |
| 10 | 7 | 0.32*** | 0.08** | 4.00 | ** |
| 11 | 34 | 0.47*** | 0.03*** | 15.67 | *** |
| 12 | 6 | 0.40*** | 0.13* | 3.08 | ** |
| 13 | 12 | 0.06ns | 0.25*** | - | - |
| 14 | 15 | 0.70*** | 0.004ns | - | - |
| 15 | 16 | 0.89*** | 0.08* | 11.13 | *** |
| 16 | 91 | 0.35*** | 0.11*** | 3.18 | *** |
| 17 | 34 | 0.27*** | 0.18*** | 1.50 | ** |
| 18 | 3 | 0.52ns | 0.14ns | - | - |
| 19 | 18 | 0.37*** | 0.13*** | 2.85 | *** |
| 20 | 42 | 0.48*** | 0.10*** | 4.80 | *** |
| 21 | 32 | 0.39*** | 0.04*** | 9.75 | *** |
| 22 | 8 | 0.09ns | 0.05ns | - | - |
| 23 | 26 | 0.33*** | 0.15*** | 2.20 | ** |
| 24 | 16 | 0.25*** | 0.20*** | 1.25 | - |
| 25 | 10 | 0.39*** | 0.11*** | 3.55 | *** |
| 26 | 17 | 0.42*** | 0.08*** | 5.25 | *** |

\*\*\*, \*\* and \* indicate significant levels at $p < 0.001$ , $p < 0.01$ , and $p < 0.05$ , and ns indicates no significance ( $p > 0.05$ ). The $Slope_{(RD-CT)}/Slope_{(RD-SR)}$ and the Slope comparison are denoted by “-” when either of the above slopes is statistically different.

**Fig. 3.**
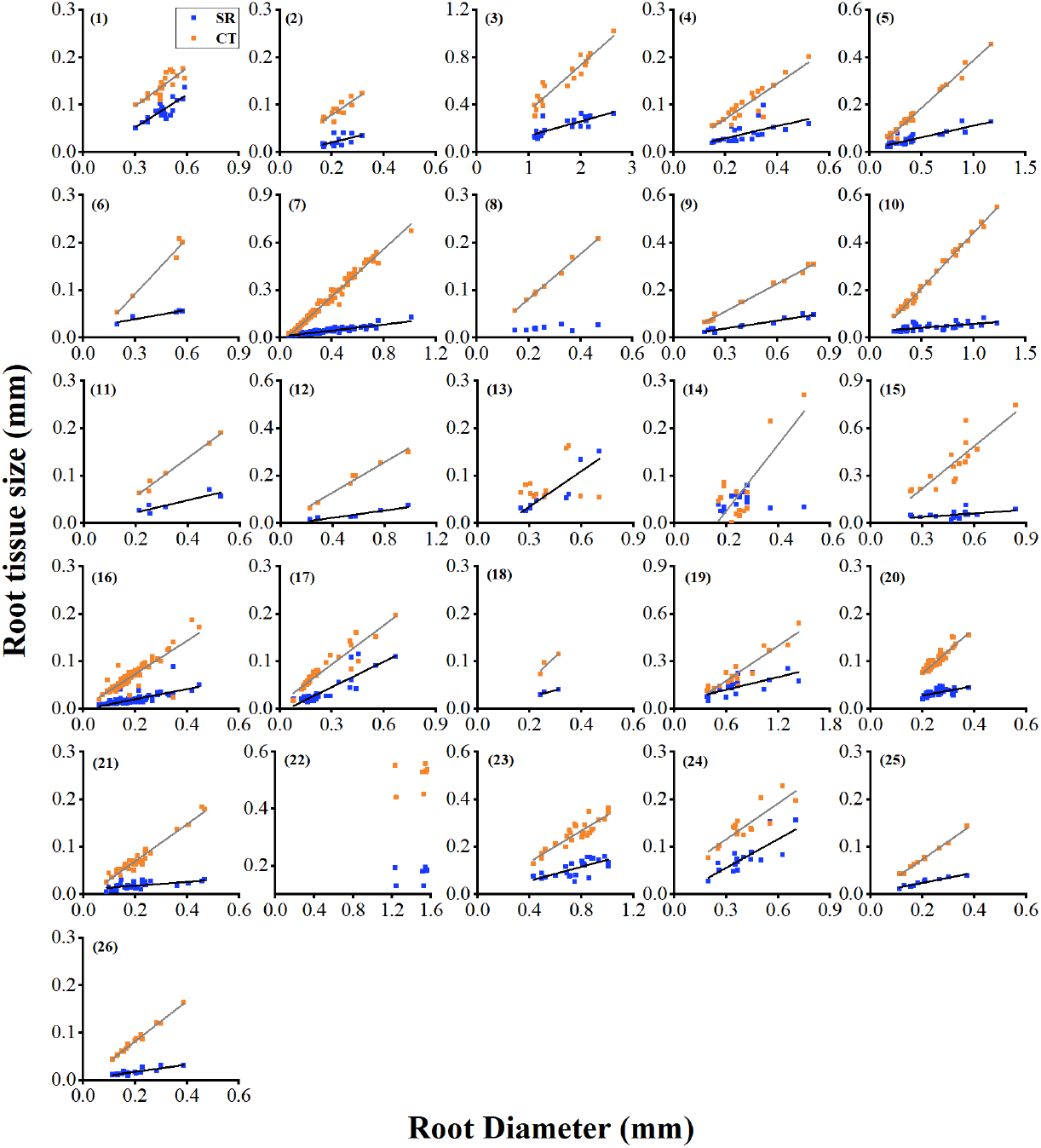
The Allometric relationship between the root cortex and the stele in 26 studies with more than three plant species examined. See supplementary data 1 and 2 for detailed information of these studies. CT: cortex thickness; SR: stele radius.

The allometric relationship in absorptive roots also occurred in different plant growth forms. For example, the thickness of tToS in woody species increases 7.6-fold faster than the stele radius with increasing root diameter, and the slope difference (3.3-fold) is much lower in non-woody species (Table 1; Fig. 4), This allometric relationship is very similar to that reported by Kong et al. (2019) using less than half of the species number as in our study Within the non-woody species, the allometric relationship was found in each of the fern grass, herb, and Orchidaceae (Fig. 5). Interestingly, we observed a 5.1-fold difference of the slope in the root allometric relationship in Orchidaceae species while the slope difference is much lower in other three non-woody groups (Table 1). Finally, for the vine species, both the woody and non-woody ones, followed the allometric manner in building their absorptive roots (Supplementary Table 1;Supplementary Fig. 2a, 2b, 2c).

**Fig. 4.**
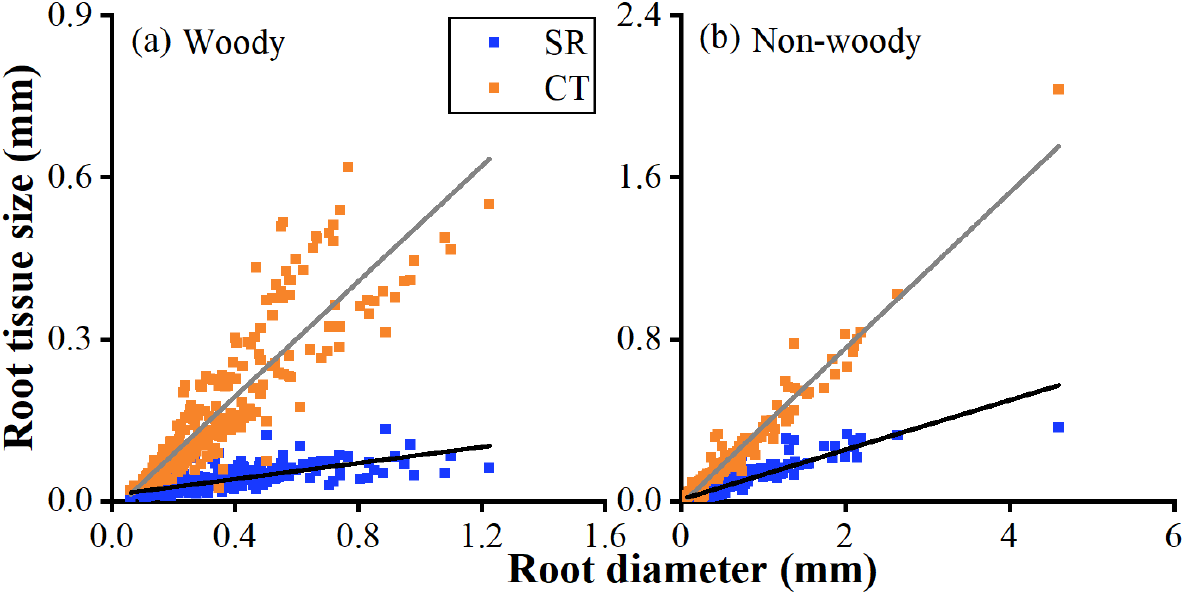
The Allometric relationship between the root cortex and the stele in woody (a) and non-woody (b) species. CT: cortex thickness; SR: stele radius.

**Fig. 5.**
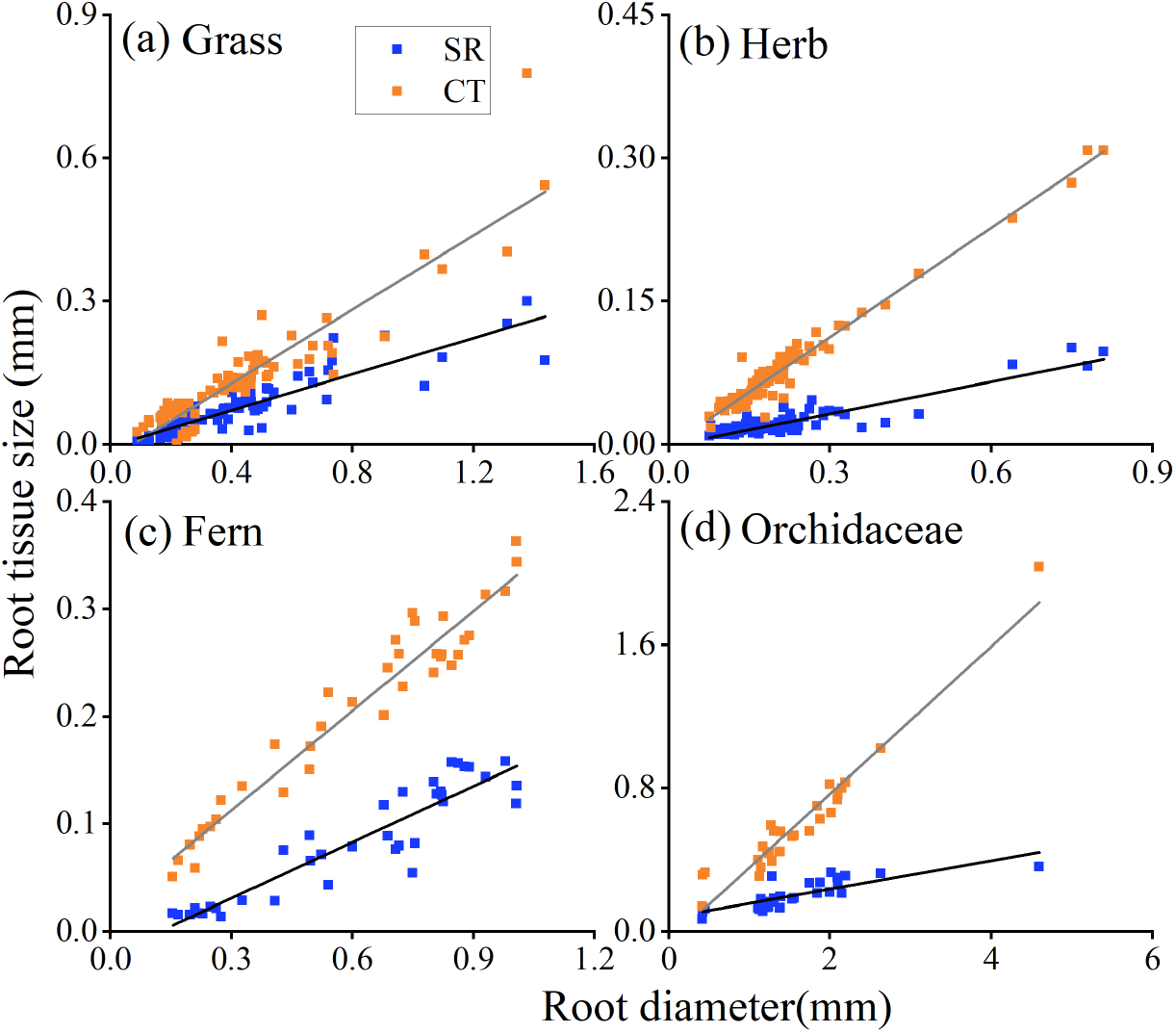
The allometric relationship between the root cortex and stele in grass (a), herb (b), fern (c) and Orchidaceae (d). CT: cortex thickness, SR: stele radius.

Among the dominant mycorrhizal types, the allometric relationship between root cortex and stele was observed in AM (Fig. 6a), regardless of being woody or non-woody of the plants (Fig. 7a, 7b), and the dual mycorrhizas of AM & EM plants but not in EM plants (Table 1; Fig. 6b, 6c). While we do note a significant allometric relationship in broadleaf EM trees but not in coniferous EM trees (Table 2; Fig. 8). Therefore, it is likely that the inclusion of such coniferous EM trees could lead to the overall no root allometric relationship across the EM plants. Considering only six coniferous EM plant species included in our dataset, we can not rule out the possibility of the root allometric relationship in other coniferous EM trees. Nevertheless, the contrasting root allometric relationships between broadleaf and coniferous EM trees may reflect the interior difference of the two types of EM plants in root structures and functioning (Guo et al., 2008b; Chen et al., 2016). the coniferous EM trees usually have vascular conduits (tracheid and sieve cells) with much lower matter (water and photosynthates) transport efficiency relative to the more efficient conduits (vessels and sieve tubes) in broadleaf EM plants (Guo et al., 2008b). It is likely that with the shift of root diameter across coniferous EM trees, some unknown strategies could be adopted rather than only change the size of the stele to increase matter transport efficiency; this, as such, results in no root allometric relationship in the coniferous EM plants. Finally, we show for the first time that the root allometric relationship still exists in another important mycorrhizal type, i.e., the Orchid mycorrhiza (OM) plants (Table 2; Fig. 5d), usually bearing much thicker absorptive roots (up to 4.6mm) (Zhu et al., 2016) than most of the AM and EM plants.

**Fig. 6.**
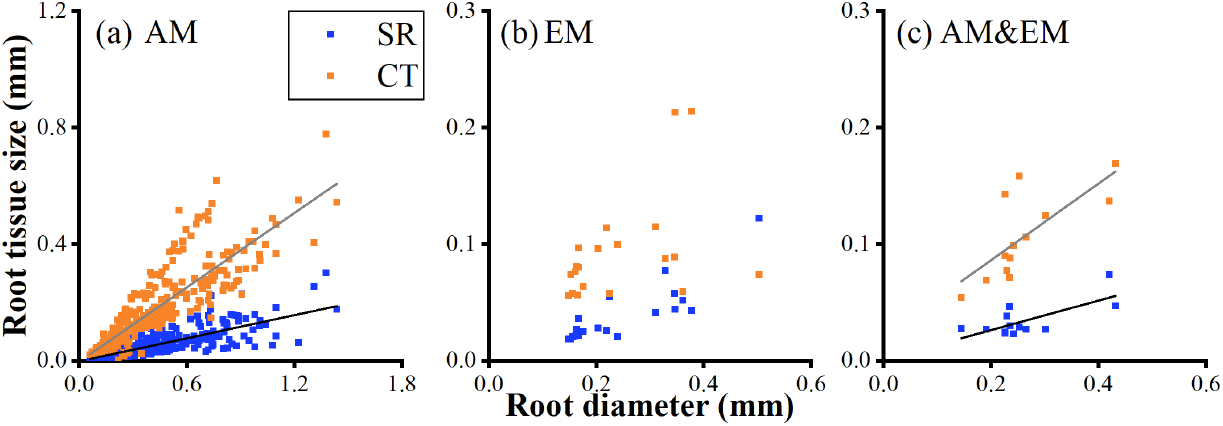
The allometric relationship between the root cortex and the stele in different mycorrhizal plants. AM: Mycorrhizae (a); EM: Ectomycorrhizae (b); AM&EM: dual mycorrhizas of AM and EM (c); CT: cortex thickness, SR: stele radius.

**Fig. 7.**
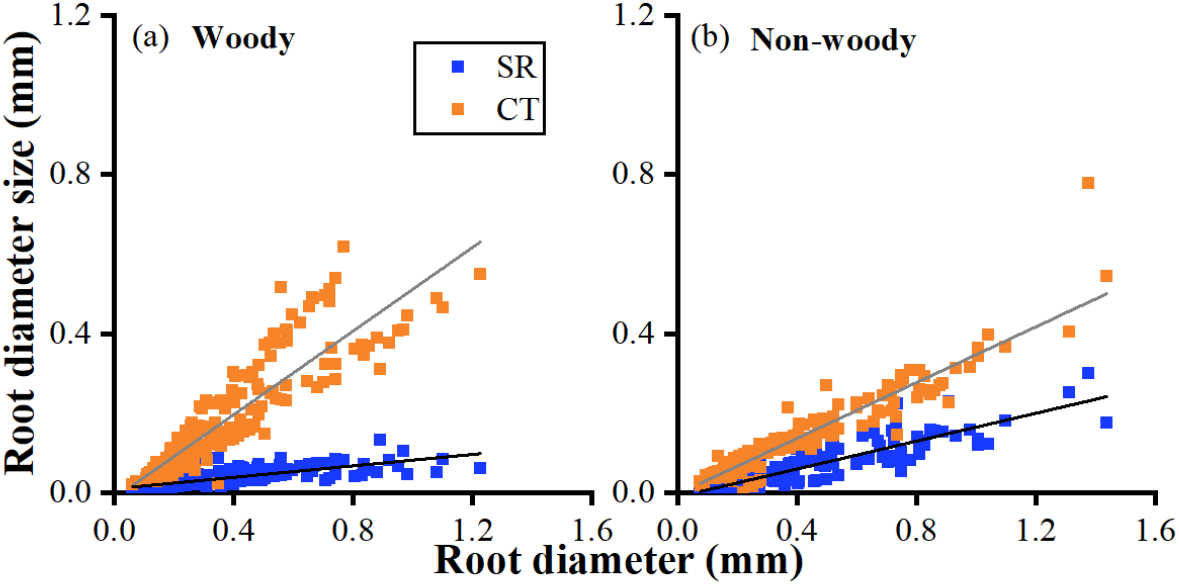
The allometric relationship between the root cortex and the stele in woody AM plants (a) and non-woody AM plants (b). CT: cortex thickness, SR: stele radius.

**Fig. 8.**
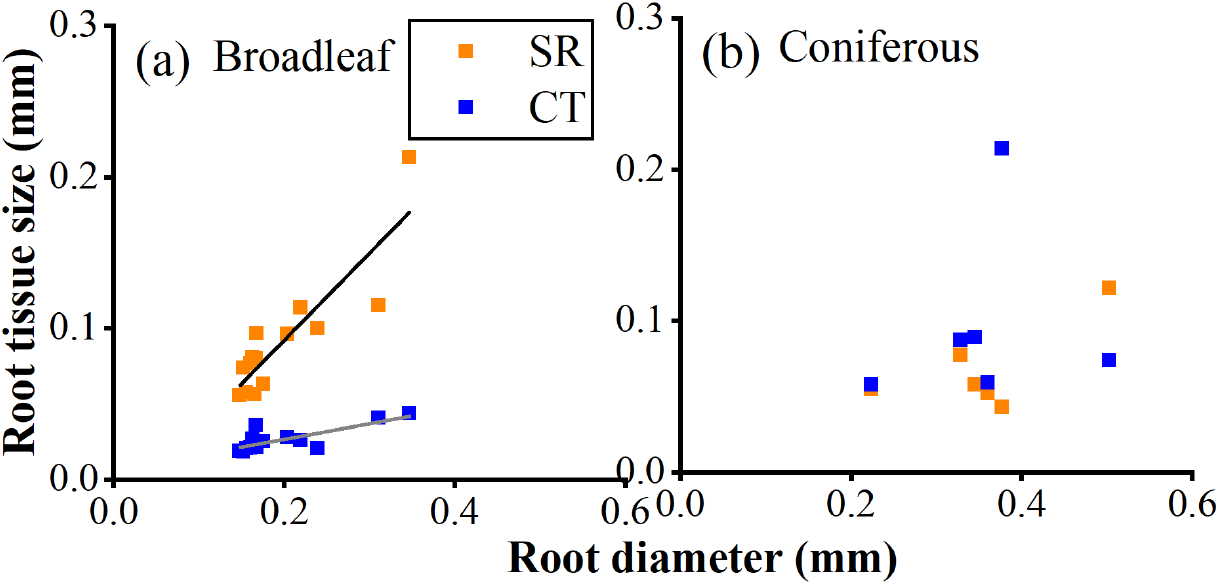
The allometric relationship between the root cortex and the stele in broadleaf EM plants (a) and coniferous EM plants (b). CT: cortex thickness, SR: stele radius.

Root anatomical structures have also been measured sparsely under different environmental change scenarios (e.g., soil nitrogen or phosphorus fertilization, increase of atmospheric CO_2_ concentration and seasonality in rainfall) (Table 1; Fig. 9), which provides an opportunity to test the consistency of the root allometric relationship. Although the root allometric relationship in some scenarios does not hold statistically (Fig. 9), this is apparently due to the inclusion of a few species with “exceptionally” large or small size of root cortex and stele or the inclusion of some species with “exceptional” responses to the environmental changes (e. g., Fig. 9a_2_, a_3_, b_2_, b_3_, d_2_, d_3_). Overall, our results suggested a relative insensitiveness of the root allometric relationship to the environmental changes. Interestingly, we note that the cortex thickness increases slower and the stele radius increases faster with increasing root diameter in the rainy season compared with that in the dry season, consequently causing an equal rather than allometric increase rate of the cortex thickness and stele radius in the rainy season (Fig. 9d_2_, d_3_). It is worthwhile to test the generality of such impact on the root allometric relationship by rainfall seasonality and uncover the underlying mechanisms in future studies.

**Fig. 9.**
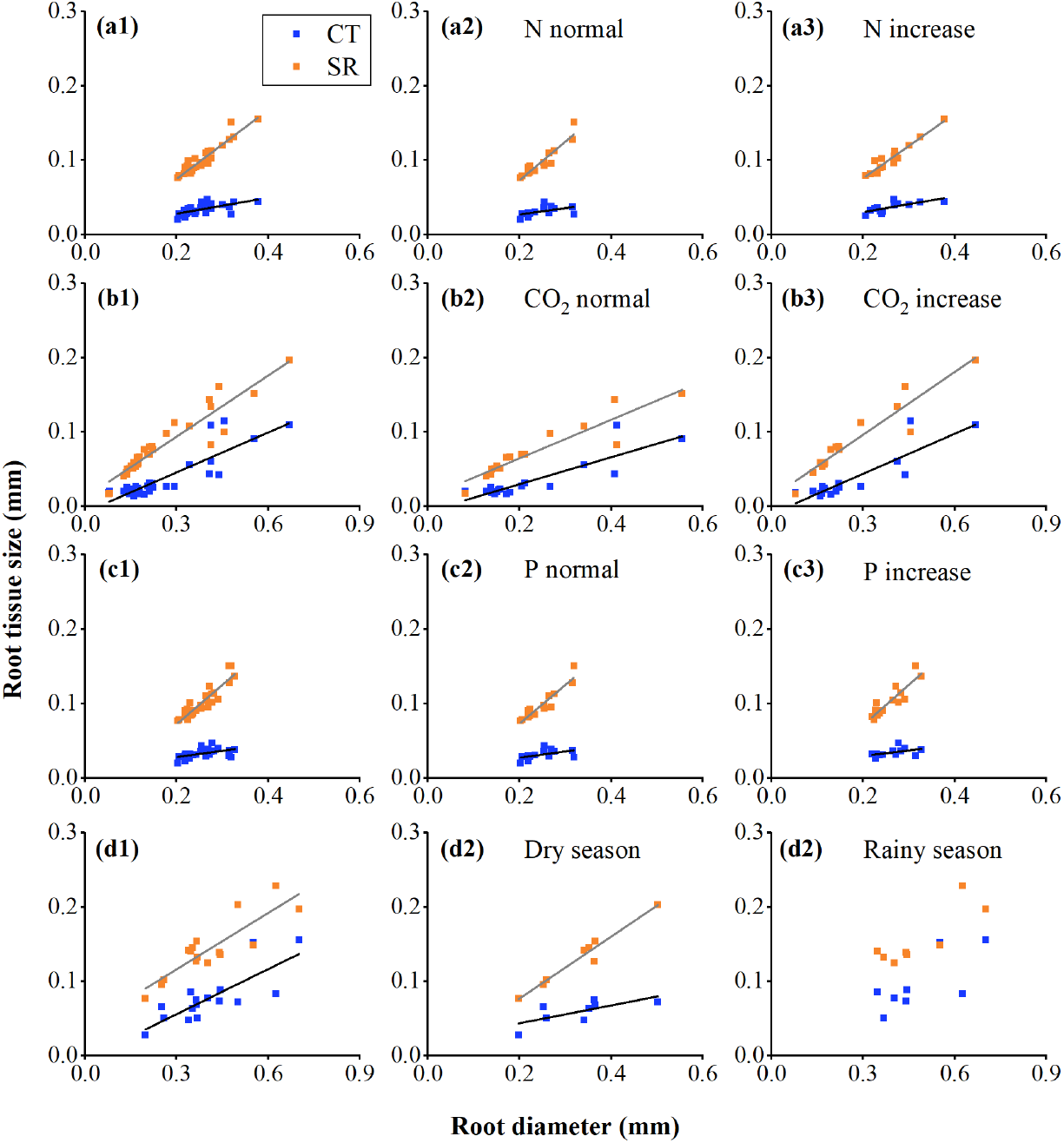
The allometric relationship between the root cortex and the stele under different environmental treatments. Nitrogen (N) deposition: control + N deposition (a_1_), control (a_2_) and N increase (a_3_); elevation of atmospheric CO_2_ concentration: control + CO_2_ increase (b_1_), control (b_2_), CO_2_ increase (b_3_); phosphorus (P) fertilization: control + P increase (c_1_), control (c_1_), P increase (c_1_); seasonality: dry season + rain season (d_1_), dry season (d_2_), rain season (d_3_). CT: cortex thickness, SR: stele radius.

Together, the wide existence of the allometric relationship between root cortex and stele across different climatic zones (tropical, sub-tropical and temperate), ecosystem types (forests, grasslands, deserts and mangroves), mycorrhizal types (AM, some EM, AM&EM, OM), phylogenetic gradients (from ferns to Orchidaceae), and environmental change scenarios support the universal rule of the allometric relationship by which the root anatomical structures are assembled.

## 3. Mechanisms of the root allometric relationship

Currently, two theories have been proposed in explaining why the allometric relationship between the cortex and stele is formed. One is the nutrient absorption-transportation balance theory (Kong et al., 2017) and the other is the carbon supply-consumption balance theory (Kong et al., 2021; Colombi et al., 2022) (Fig. 10). Both theories run according to the principle of functional balance of the matter (nutrients, photosynthates) transport within root tissues as well as the physical law of fluid transport in the conduits, namely the “Hagen-Poiseuille law” (Jensen et al., 2016). Here, we only outline the two theories and readers can refer to the original papers for details of the theories.

**Fig. 10.**
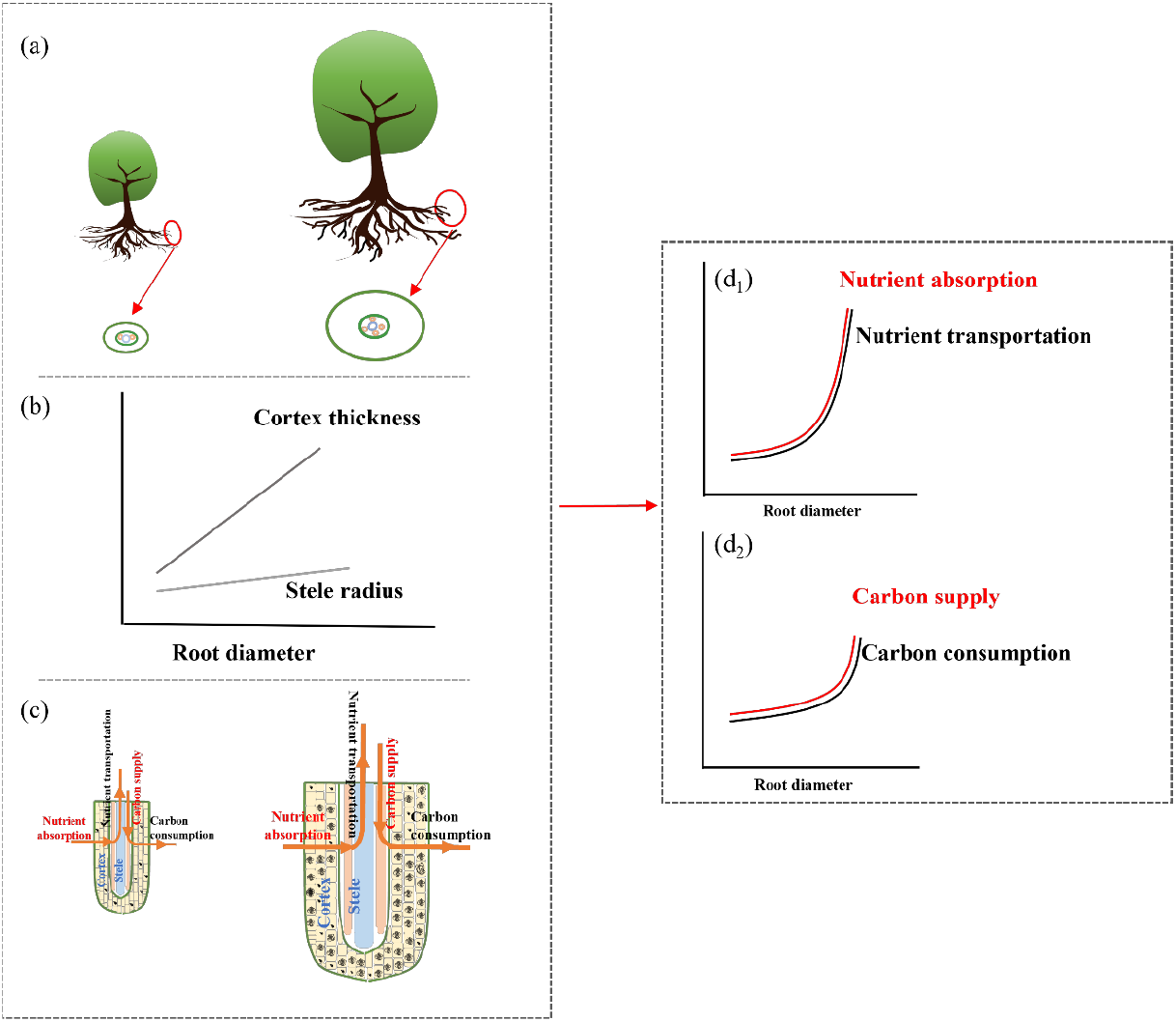
Mechanisms for the allometric relationship between the cortex and the stele in absorptive roots. For simplicity, we presented two plant species with a thin and a thick absorptive root, respectively, followed by the cross-sectional area of the roots (a). The cortex section is indicated by the area between the two green circles; the inner green circle, the stele; blue circles, the vessels; orange circles, the sieves. Different functions of the root anatomical structures are shown in the longitudinal section model of the roots (b)The change in the size of the cortex and stele with the shift of root diameter across species is shown in (c): cortex, sites for the symbiosis with mycorrhizal fungi (i.e., the intermingled lines in the cortical cells) and carbon consumption; vessels, transport for the water and nutrients; sieves, transport for carbohydrates. The allometric relationship between the root cortex and stele is considered to meet the balance between nutrient absorption (via mycorrhizal fungi in the cortex) and nutrient transportation (via the stele) (d_1_) and between carbon supply (via the stele) and carbon consumption (via the cortex) (d_2_). The conceptual models in (d_1_) and (d_2_) are redrawn from Kong et al. (2017) and Kong et al. (2021).

There are two parallel vascular systems in root steles, i.e., vessels responsible for transporting water and nutrients upward to stems and leaves and sieves responsible for meeting the carbon demanding of the root. According to the Hagen-Poiseuille law, both volumetric flow rates in the conduits (i.e., water and nutrient transportation via vessels and photosynthate transportation via sieves) scale to the fourth power of the root radius; while even the maximum nutrient absorption (via mycorrhizal fungi in the cortex) and carbon consumption (via the cortex) scale less than the twice power of root radius. In this case, only a much faster increase of the cortex thickness than the stele radius with increasing root diameter (i.e., the allometric relationship) can lead to a *balance between the nutrient absorption* (via mycorrhizal fungi in the cortex) *and the nutrient transportation* (via vessels in the stele) and *a balance between the carbon supply* (via sieves in the stele) *and carbon consumption* (via the cortex).

Nevertheless, we should also keep in mind of some important limitations of the above two theories. Firstly, the *nutrient absorption-transportation balance* theory holds on a prerequisite of a universal association of plant roots with mycorrhizal fungi, while there are still a lot of species with no mycorrhizal associations (Vander et al., 2015; Brundrett and Tedersoo, 2018; Correia et al., 2018). Secondly, the two theories seem running independently although they are based on two interconnected vascular conducts, i.e., vessels and sieves. It is also interesting to learn about how the above two theories are linked with leaves, the important sink of nutrients and the source of carbon. Thirdly, empirical evidence is urgently needed to test the prediction of the above functional balance that underlies the root allometric relationship.

## 4. Implication of the root allometric relationship

### 4.1 Relationship with the “root economics spectrum”

Traditionally, root economic spectrum is considered to be the core trait dimension in roots which conveys a trade-off between nutrient uptake and conservation (Freschet et al., 2010; Reich, 2014; Kramer-Walter et al., 2016). It has been widely recognized that a positive correlation between root diameter and root life span (i.e., conservation of nutrient) (Guo et al., 2008a; Gu et al., 2017; Liese et al., 2019) and a negative correlation between root nutrient uptake and root tissue density (RTD) (Zadworny et al., 2017; Stock et al., 2021). If the trade-off exists between root nutrient uptake and conservation, there should be a positive correlation between RTD and root diameter. However, besides such a prediction, many studies also found a negative or no relationship between RTD and root diameter (Weemstra et al., 2016; Kong et al., 2019; Han and Zhu, 2021). Based on the universal allometric relationship between root cortex and stele, we predict a negative and non-linear relationship between RTD and root diameter, and this prediction has been verified using a global root trait dataset (Kong et al., 2019). Therefore, the formation of the root allometric relationship may not support the existence of the widely acknowledged root economic spectrum.

### 4.2 Plant evolution and adaptation to environment

The evolution of angiosperms is closely related to the decline of atmospheric CO_2_ concentration since the mid-Cretaceous (Beerling and Berner, 2005; Gerhart and Ward, 2010). For example, the reduction of atmospheric CO_2_ concentration often lowers leaf photosynthesis, consequently causing “carbon starvation” to plants. To survive in this carbon limitation condition, plants tend to increase stomatal conductance to compensatively improve leaf CO_2_ fixation (Zhou et al., 2013; Holtta et al., 2017). However, large stomatal conductance will enhance the transpiration water loss, as such causing physiological drought to plants (Khan et al., 2007; Wang et al., 2018). The surge of leaf vein density in angiosperms since the mid-Cretaceous, hence resulting more efficient water supply to the mesophyll cell for photosynthesis, can be considered as evidence for plant adaptation to the physiological drought (Baraloto et al., 2010; Feild et al., 2011; Baird et al., 2021; Yan et al., 2022).

Coordinated with the evolutionary change in leaves, thinning of the absorptive roots is regarded as an adaptation to the physiological drought caused by the decline of atmospheric CO_2_ concentration (Fig. 11a) (Comas et al., 2012; Chen et al., 2013; Ma et al., 2018). Alongside the thinning of the absorptive roots, the much faster decrease of the cortex thickness than the stele radius means effectively reducing the resistance of water and nutrients entering the root tissues for one hand and for the other hand effectively reducing the carbon consumption by root cortex (Fig. 11b). Therefore, the allometrically structured roots are much beneficial for plants to survive under the carbon and water limited environment. From this point of view, the allometric relationship in absorptive roots is insightful for our understanding of how the roots, whole plants and even the ecosystems respond and adapt to the geological and the on-going environmental changes.

**Fig. 11.**
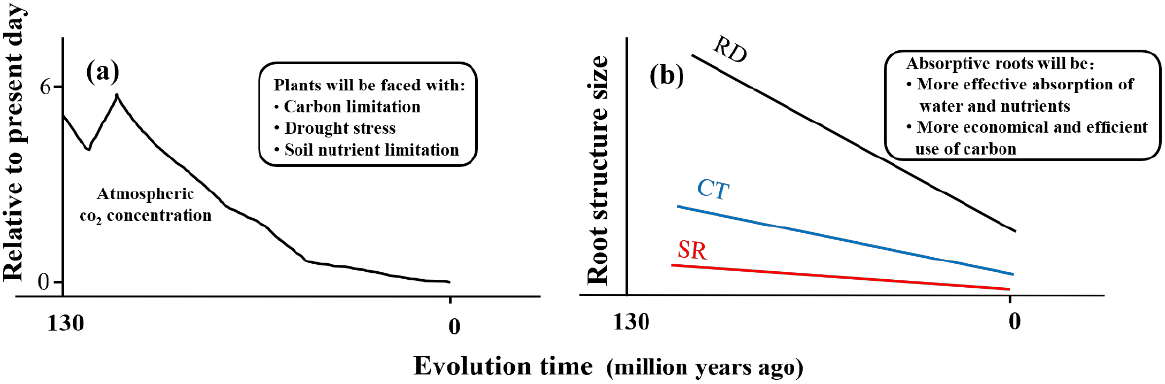
The co-variation of atmospheric CO_2_ concentration (a) and anatomical structures in absorptive roots (b) since the Cretaceous. The pattern for the change of atmospheric CO_2_ concentration is redrawn from the study by Beerling et al. (2010). The resulting environmental changes and the adaptive responses of the roots are presented as inlets in this figure. RD: absorptive root diameter; CT: cortex thickness, SR: stele radius.

## 5. Future directions

### 5.1 Differences of the root allometric relationship among studies

Studies to date always concentrate on the general pattern of the root allometric relationship, i.e., a much larger slope of the cortex thickness *vs*. root diameter regression than the slope of the stele radius *vs*. root diameter regression, while ignore the great difference of the above allometry, that is, the slope difference ranging from the minimum 1.2-fold to the maximum 15.7-fold across studies (Table 2). We are still unclear about mechanisms accounting for such huge difference in the root allometric relationship. This is a fascinating question that could stimulate far-reaching outcomes in this filed.

### 5.2 Examining root anatomical structures in more species

The global establishment of the allometric relationship between root cortex and stele relies on about 500 plant species, much smaller than the total vascular plant species number (about 390,000) on the earth (Cantwell-Jones et al., 2022). Even in the global root trait dataset, such as FRED 3.0 (Iversen et al., 2021), the measurements of root anatomical traits are far less than the measurements of other root traits such as root diameter and root tissue density. Therefore, it is necessary to measure root anatomical structures in more plant species, especially the families with a large number of species like Orchidaceae with over 20, 000 species.

### 5.3 Effects of environmental changes on the root allometric relationship

By far, only four studies are available for our evaluation on how environmental changes alter the allometric relationship in absorptive roots. Moreover, only a few environmental change scenarios are considered and no interactions among these factors have been examined in these studies. Apart from the studies under controlled environments, we need to pay more attention to plants growing in naturally stressful conditions such as alpine forests, deserts, and coast environments in high salinity. Comparison of the root allometric relationship under the controlled and the natural conditions could be instructive for our understanding and prediction of vegetation dynamics under global climate change.

### 5.4 Linking the root allometric relationship with plant above-ground organs

Plant growth and evolution depends on the functional coordination between plant above- and below-ground organs (Aritsara et al., 2022; Weigelt et al., 2021; Zhou et al., 2022). In the framework of the root allometric relationship, the faster increase of the cortex thickness than stele radius could be accompanied with a faster supply of leaf photosynthate to the roots. We know little about how plants with thick absorptive roots assign their leaf traits to meet functional balance of water, nutrients and carbon between roots and leaves. Another interesting question is how the root allometric relationship can be coordinated with plant reproductive organs like flowers, fruits and seeds given that the reproductive organs usually compete with roots for photosynthates and with leaves for water and nutrients. Therefore, linking the root allometric relationship with plant above-ground organs could pave a new way for our understanding of the co-evolution within plants and between plants and animals for pollination or seed dispersal.

## Author Contributions

Y.Z. and D.K. conceived the ideas of this review paper, D.K. and Y.Z. performed the data analysis, Y.Z. wrote the first draft of this manuscript and Y.Z., D.K., J.C., Q.Y., M.W., and Y.Z. all contributed to the editing and revision of the final version of the manuscript.

## Acknowledgements

We are grateful for Oscar J. Valverde-Barrantes and WenHua Xiang for their assistance in data collection work. This study was funded by the National Natural Science Foundation of China (32171746, 42077450, 31870522 and 31670550), and the Scientific Research Foundation of Henan Agricultural University (30500854), Research Funds for overseas returnee in Henan Province, China.

## References

Aritsara, A.N.A., Wang, S., Li, B.N., et al., 2022. Divergent leaf and fine root “pressure-volume relationships” across habitats with varying water availability. Plant Physiol. 2–14.

Baird, A.S., Taylor, S.H., Pasquet-Kok, J., et al., 2021. Developmental and biophysical determinants of grass leaf size worldwide. Nature. 592, 242–+.

Baraloto, C., Paine, C.E.T., Poorter, L., et al., 2010. Decoupled leaf and stem economics in rain forest trees. Ecol Lett. 13, 1338–1347.

Beerling, D.J., Berner, R.A., 2005. Feedbacks and the coevolution of plants and atmospheric CO_2_. Proc Natl Sci. U. S.A. 102, 1302–1305.

Beerling, D.J., Franks, P.J., 2010. The hidden cost of transpiration. Nature. 464, 495–496.

Bergmann, J., Weigelt, A., van Der Plas. F., et al., 2020. The fungal collaboration gradient dominates the root economics space in plants. Sci Adv. 6, 1193–1208.

Brundrett, M.C., 2002. Coevolution of roots and mycorrhizas of land plants. New Phytol. 154, 275–304.

Brundrett, M.C., Tedersoo, L., 2018. Evolutionary history of mycorrhizal symbioses and global host plant diversity. New Phytol. 220, 1108–1115.

Cantwell-Jones, A., Ball, J., Collar, D., et al., 2022. Global plant diversity as a reservoir of micronutrients for humanity. Nat Plants. 8, 225–+.

Carmona, C.P., Bueno, C.G., Toussaint, A., et al., 2021. Fine-root traits in the global spectrum of plant form and function. Nature. 597, 683–687.

Chandregowda, M.H., Tjoelker, M.G., Pendall, E., et al., 2022. Root trait shifts towards an avoidance strategy promote productivity and recovery in C-3 and C-4 pasture grasses under drought. Funct Ecol. 36, 1754–1771.

Chen, W.L., Koide, R.T., Adams, T.S., et al., 2016. Root morphology and mycorrhizal symbioses together shape nutrient foraging strategies of temperate trees. Proc Natl Sci. U. S.A. 113, 8741–8746.

Chen, W.L., Zeng, H., Eissenstat, D.M., et al., 2013. Variation of first-order root traits across climatic gradients and evolutionary trends in geological time. Glob Ecol Biogeogr. 22, 846–856.

Iversen C.M., McCormack M.L., Baer J.K., et al., 2021. Fine-Root Ecology Database (FRED): A Global Collection of Root Trait Data with Coincident Site, Vegetation, Edaphic, and Climatic Data, Version 3.

Colombi, T., Chakrawal, A., Herrmann, A.M., 2022. Carbon supply-consumption balance in plant roots: effects of carbon use efficiency and root anatomical plasticity. New Phytol. 233, 1542–1547.

Comas, L.H., Mueller, K.E., Taylor, L.L., et al., 2012. Evolutionary patterns and biogeochemical significance of angiosperm toot traits. Int J Plant Sci. 173, 584–595.

Correia, M., Heleno, R., Vargas, P., et al., 2018. Should I stay or should I go? Mycorrhizal plants are more likely to invest in long-distance seed dispersal than non-mycorrhizal plants. Ecol Lett. 21, 683–691.

Diaz, S., Kattge, J., Cornelissen, J.H.C., et al., 2016. The global spectrum of plant form and function. Nature. 529, 167–+.

Eissenstat, D.M., Kucharski, J.M., Zadworny, M., et al., 2015. Linking root traits to nutrient foraging in arbuscular mycorrhizal trees in a temperate forest. New Phytol. 208, 114–124.

Encinas-Valero, M., Esteban, R., Hereş, A.M., et al., 2022. Holm oak decline is determined by shifts in fine root phenotypic plasticity in response to belowground stress. New Phytol. 235, 2237–2251.

Feild, T.S., Brodribb, T.J., Iglesias, A., et al., 2011. Fossil evidence for Cretaceous escalation in angiosperm leaf vein evolution. Proc Natl Sci. U. S.A. 108, 8363–8366.

Freschet, G.T., Cornelissen, J.H.C., Van Logtestijn, R.S.P., et al., 2010. Evidence of the ‘plant economics spectrum’ in a subarctic flora. J Ecol. 98, 362–373.

Gerhart, L.M., Ward, J.K., 2010. Plant responses to low CO_2_ of the past. New Phytol. 188, 674–695.

Gu, J.C., Wang, Y., Fahey, T.J., et al., 2017. Effects of root diameter, branch order, soil depth and season of birth on fine root life span in five temperate tree species. Eur J Forest Res. 136, 727–738.

Gu, J.C., Xu, Y., Dong, X.Y., et al., 2014. Root diameter variations explained by anatomy and phylogeny of 50 tropical and temperate tree species. Tree Physiol. 34, 415–425.

Guo, D.L., Mitchell, R.J., Withington, J.M., et al., 2008a. Endogenous and exogenous controls of root life span, mortality and nitrogen flux in a longleaf pine forest: root branch order predominates. J Ecol. 96, 737–745.

Guo, D.L., Xia, M.X., Wei, X., et al., 2008b. Anatomical traits associated with absorption and mycorrhizal colonization are linked to root branch order in twenty-three Chinese temperate tree species. New Phytol. 180, 673–683.

Han, M.G., Zhu, B., 2021. Linking root respiration to chemistry and morphology across species. Glob Change Biol. 27, 190–201.

Holtta, T., Lintunen, A., Chan, T., et al., 2017. A steady-state stomatal model of balanced leaf gas exchange, hydraulics and maximal source-sink flux. Tree Physiol. 37, 851–868.

Hong, Y., Zhou, Q., Hao, Y., et al., 2022. Crafting the plant root metabolome for improved microbe-assisted stress resilience. New Phytol. 234, 1945–1950.

Jensen, K. H., Berg-Sørensen, K., Bruus, H., et al., 2016. Sap flow and sugar transport in plants. Rev Mod Phys. 88, 035007.

Joswig, J.S., Wirth, C., Schuman, M.C., et al., 2022. Climatic and soil factors explain the two-dimensional spectrum of global plant trait variation. Nat Ecol Evol. 6, 36–+.

Khan, H.U.R., Link, W., Hocking, T.J., et al., 2007. Evaluation of physiological traits for improving drought tolerance in faba bean (Vicia faba L.). Plant Soil. 292, 205–217.

Kong, DL., Wang, J., Zeng, H., et al., 2014. Leading dimensions in absorptive root trait variation across 96 subtropical forest species. New Phytol. 203, 863–872.

Kong, DL., Wang, J., Wu, H., et al., 2019. Nonlinearity of root trait relationships and the root economics spectrum. Nat Commun. 10, 2203.

Kong, DL., Wang, J., Zeng, H., et al., 2017. The nutrient absorption–transportation hypothesis: optimizing structural traits in absorptive roots. New Phytol. 213, 1569–1572.

Kong, DL., Wang, J.J., Valverde-Barrantes, O.J., et al., 2021. A framework to assess the carbon supply-consumption balance in plant roots. New Phytol. 229, 659–664.

Kramer-Walter, K.R., Bellingham, P.J., Millar, T.R., et al., 2016. Root traits are multidimensional: specific root length is independent from root tissue density and the plant economic spectrum. J Ecol. 104, 1299–1310.

Laughlin, D.C., Mommer, L., Sabatini, F.M., et al., 2021. Root traits explain plant species distributions along climatic gradients yet challenge the nature of ecological trade-offs. Nat Ecol Evol. 5, 1123–+.

Li, X., Dong, J.L., Chu, W.Y., et al., 2018. The relationship between root exudation properties and root morphological traits of cucumber grown under different nitrogen supplies and atmospheric CO_2_ concentrations. Plant Soil. 425, 415–432.

Liese, R., Leuschner, C., Meier, I.C., 2019. The effect of drought and season on root life span in temperate arbuscular mycorrhizal and ectomycorrhizal tree species. J Ecol. 107, 2226–2239.

Lugli, L.F., Rosa, J.S., Andersen, K.M., et al., 2021. Rapid responses of root traits and productivity to phosphorus and cation additions in a tropical lowland forest in Amazonia. New Phytol. 230, 116–128.

Ma, ZQ., Guo, DL., Xu, X., et al., 2018. Evolutionary history resolves global organization of root functional traits. Nature. 555, 94–97.

Martin, F.M., Uroz, S., Barker, D.G., 2017. Ancestral alliances: Plant mutualistic symbioses with fungi and bacteria. Science. 356, eaad4501

McCormack, M.L., Guo, D.L., Iversen, C.M., et al., 2017. Building a better foundation: improving root-trait measurements to understand and model plant and ecosystem processes. New Phytol. 215, 27–37.

Pineiro, J., Ochoa-Hueso, R., Drake, J.E., et al., 2020. Water availability drives fine root dynamics in aEucalyptuswoodland under elevated atmospheric CO(2)concentration. Funct Ecol. 34, 2389–2402.

Reich, P.B., 2014. The world-wide ‘fast–slow’ plant economics spectrum: a traits manifesto. J Ecol. 102, 275–301.

Rich, M.K., Vigneron, N., Libourel, C., et al., 2021. Lipid exchanges drove the evolution of mutualism during plant terrestrialization. Science. 372, 864–+.

Stock, S.C., Koester, M., Boy, J., et al., 2021. Plant carbon investment in fine roots and arbuscular mycorrhizal fungi: A cross-biome study on nutrient acquisition strategies. Sci Total Environ. 781, 146748.

Van der Heijden, M.G.A., Martin, F.M., Selosse, M.A., et al., 2015. Mycorrhizal ecology and evolution: the past, the present, and the future. New Phytol. 205, 1406–1423.

Wambsganss, J., Freschet, G.T., Beyer, F., et al., 2021. Tree species mixing causes a shift in fine-root soil exploitation strategies across European forests. Funct Ecol. 35, 1886–1902.

Wang, X.X., Du, T.T., Huang, J.L., et al., 2018. Leaf hydraulic vulnerability triggers the decline in stomatal and mesophyll conductance during drought in rice. J Exp Bot. 69, 4033–4045.

Weemstra, M., Mommer, L., Visser, E.J., et al., 2016. Towards a multidimensional root trait framework: a tree root review. New Phytol. 211, 1159–1169.

Weigelt, A., Mommer, L., Andraczek, K., et al., 2021. An integrated framework of plant form and function: the belowground perspective. New Phytol. 232, 42–59.

Wen, Z.H., White, P.J., Shen, J.B., et al., 2022. Linking root exudation to belowground economic traits for resource acquisition. New Phytol. 233, 1620–1635.

Wright, I.J., Reich, P.B., Westoby, M., et al., 2004. The worldwide leaf economics spectrum. Nature. 428, 821–827.

Yan, H., Freschet, G.T., Wang, H.M., et al., 2022. Mycorrhizal symbiosis pathway and edaphic fertility frame root economics space among tree species. New Phytol. 234, 1639–1653.

Zadworny, M., McCormack, M.L., Zytkowiak, R., et al., 2017. Patterns of structural and defense investments in fine roots of Scots pine (Pinus sylvestris L.) across a strong temperature and latitudinal gradient in Europe. Glob Change Biol. 23, 1218–1231.

Zhou, M., Guo, Y.M., Sheng, J., et al., 2022. Using anatomical traits to understand root functions across root orders of herbaceous species in a temperate steppe. New Phytol. 234, 422–434.

Zhou, Y.M., Jiang, X.J., Schaub, M., et al., Ten-year exposure to elevated CO_2_ increases stomatal number of Pinus koraiensis and P. sylvestriformis needles. Eur J Forest Res. 132, 899–908.

Zhu L.Q., Xu Y.X., Zhao L.J., et al., 2016. anatomical structure and environmental adaptability of Cymbidium cyperifolium in karst area. Guihaia. 36, 1179–1185+1164.

